# Intracranial self-stimulation and concomitant behaviors following systemic methamphetamine administration in *Hnrnph1* mutant mice

**DOI:** 10.1101/2020.06.05.137190

**Authors:** Kristyn N. Borrelli, Carly R. Langan, Kyra R. Dubinsky, Karen K. Szumlinski, William A. Carlezon, Elena H. Chartoff, Camron D. Bryant

## Abstract

**Rationale:** Addiction to methamphetamine (MA) is a major public health issue in the United States. While psychostimulant use disorders are heritable, their genetic basis remains poorly understood. We previously identified heterogeneous nuclear ribonucleoprotein H1 (*Hnrnph1;* H1) as a quantitative trait gene underlying sensitivity to MA-induced locomotor activity. Mice heterozygous for a frameshift deletion in the first coding exon of H1 (H1^+/-^) showed reduced MA phenotypes including oral self-administration, locomotor activity, dopamine release, and dose-dependent differences in MA conditioned place preference. However, the effects of H1^+/-^ on innate and MA-modulated reward sensitivity are not known.

**Objectives:** We examined innate reward sensitivity and modulation by MA in H1^+/-^ mice via intracranial self-stimulation (ICSS).

**Methods:** We used intracranial self-stimulation (ICSS) of the medial forebrain bundle to assess shifts in reward sensitivity following acute, ascending doses of MA (0.5-4.0 mg/kg, i.p.) using a within-subjects design. We also assessed video-recorded behaviors during ICSS testing sessions.

**Results:** H1^+/-^ mice displayed reduced normalized maximum response rates, H1^+/-^ females showed lower normalized M50 values compared to wild-type females following MA, and H1^+/-^ influenced ICSS responding relative to maximum baseline rates. There was a dose-dependent reduction in distance to the response wheel following MA administration, providing an additional measure of reward-related behavior.

**Conclusions:** H1^+/-^ mice displayed reduced reward facilitation following MA in a sex- and dose-dependent manner. This result expands upon the set of MA-induced phenotypes observed in H1^+/-^ mice.

## INTRODUCTION

Abuse of psychostimulant drugs, such as methamphetamine (**MA**), contributes to an ongoing public health crisis marked by unprecedented overdose rates in the United States. Deaths linked to psychostimulant misuse increased by 33.3 percent from 2016-2017 alone (Seth et al. 2018). Psychostimulant use disorder has a significant heritable component, but the genetic basis remains poorly understood (Bousman et al. 2009a; Jensen 2016; Goldman et al. 2005). Prior clinical studies designed to discover genes associated with MA addiction traits have been hindered by several limitations, including a lack of statistical power and environmental control (Uhl et al. 2008; Ikeda et al. 2013; Hancock et al. 2018). Rodent models help address some of these impediments and permit the study of both the genetic and neurobiological mechanisms underlying addiction-relevant quantitative traits. Forward genetic approaches, such as quantitative trait locus (**QTL**) mapping, can identify chromosomal loci containing novel genes and variants linked to addiction-relevant behaviors (Flint et al. 2005; Spanagel 2013).

We recently used QTL mapping, positional cloning, and gene editing to validate *Hnrnph1* (heterogeneous nuclear ribonucleoprotein H1; **H1**) as a quantitative trait gene underlying sensitivity to MA-induced locomotor stimulation. To validate the quantitative trait gene within the identified 0.2 Mb locus, we used transcription activator-like effector nucleases (**TALENs**) to induce a frameshift deletion in the first coding exon of the H1 gene in B6 mice. H1 mutant mice heterozygous for the *H1* deletion (**H1**^**+/-**^) recapitulated the decreased sensitivity to MA-induced locomotor activity observed in the congenic lines (Yazdani et al. 2015). Additionally, striatal transcriptomic analysis of H1^+/-^ mice suggested deficient development of dopaminergic (DA-ergic) innervation and transmission in the mesocorticolimbic reward pathway (Yazdani et al. 2015). More recently, we showed that a 0.1 Mb locus containing *Hnrnph1* was also sufficient to reduce MA behavior and was linked to decreased 5’ UTR usage of *Hnrnph1* and decreased hnRNP H protein (Ruan et al. 2020a).

*H1* codes for an RNA-binding protein that is ubiquitously expressed throughout the mouse brain (Lein et al. 2007) and regulates multiple stages of RNA metabolism, including alternative splicing (Kim et al. 2002; Han et al. 2010). Accumulating evidence indicates a crucial role for RNA-binding proteins, such as H1, in neurobehavioral plasticity underlying substance use disorders (Bryant and Yazdani, 2016). Proteomic analysis of synaptosomal striatal tissue at 30 min following an acute dose of MA (2 mg/kg, i.p.) indicated that the molecular mechanisms underlying the MA-induced behavioral deficits in H1^+/-^ mice could involve mitochondrial dysfunction as opposing effects of MA on the levels of several synaptosomally localized mitochondrial proteins were observed as a function of genotype and as a function of MA treatment (Ruan et al. 2020b).

Intracranial self-stimulation (**ICSS**) can assess innate reward sensitivity and the abuse potential and motivational effects of various addictive substances, including psychostimulants such as MA (Bauer et al. 2013; Carlezon and Chartoff 2007; Kesby et al. 2018; Negus and Miller 2014). One version of the ICSS procedure involves indirect electrical stimulation of the mesolimbic dopaminergic reward pathway by targeting the medial forebrain bundle (**MFB**); a collection of axonal fibers of passage in the lateral hypothalamus (Bielajew and Shizgal 1986; Garris et al. 1999). Stimulation of several brain regions can support ICSS; within the MFB, stimulation activates mesolimbic dopaminergic projections from the ventral tegmental area (**VTA**) to the nucleus accumbens (**NAc**) to increase synaptic dopamine levels within the NAc (Fiorino et al. 1993; Miliaressis et al. 1991; Owesson-White et al. 2008), a brain region critical for endogenous reward signaling (Schultz 2000). MA also targets plasma membrane dopamine transporters and the vesicular monoamine transporters (Cruickshank and Dyer 2009) to increase synaptic dopamine in the NAc (Johnson et al. 2018; Sulzer et al. 2005) that tracks closely with an increase in locomotor activity (Koshikawa et al. 1989; Di Chiara and Imperato 1988; Zocchi et al. 1997). The rise in NAc DA levels elicited by amphetamine-type psychostimulants can increase ICSS response rates and sensitivity to brain stimulation reward in rodent models. Conversely, drugs that deplete or inhibit DA transmission decrease sensitivity to brain stimulation reward, an indicator of anhedonia (Negus and Miller 2014; Bauer et al. 2014; Bauer et al. 2013; Phillips et al. 1989; Riday et al. 2012).

*In vivo* microdialysis studies have shown that **H1**^**+/-**^ mice have significantly lower MA-induced extracellular DA levels in the NAc compared to wild-type (**WT**) littermates following intraperitoneal (i.p.) injections of 0.5 and 2.0 mg/kg MA, in the absence of a significant difference in baseline total or extracellular DA levels (Ruan et al. 2020b). We examined the effects of H1^+/-^ on ICSS responding under baseline conditions and following “priming” of the reward circuitry with increasing doses of MA (0.5, 1.0, 2.0, 4.0 mg/kg). Because drug-induced DA release influences both ICSS and locomotor activity and because H1^+/-^ mice showed a robust reduction in MA-induced locomotor activity (Yazdani et al. 2015; Ruan et al. 2020b), we hypothesized that the threshold-reducing effects of MA would be less robust in H1^+/-^ mice compared to WT mice. We also assessed concomitant behavioral activity and location during ICSS testing sessions to potentially identify unique and potentially genotype-specific behaviors altered by MA treatment that coincide with MA-induced changes in ICSS operant responding.

## METHODS

### Mice

All procedures in mice were approved by the Boston University Institutional Animal Care and Use Committee (AN-15607; PROTO201800421) and were conducted in strict accordance with National Institute of Health Guidelines for the Care and Use of Laboratory Animals (National Research Council 2011). Colony rooms were maintained on a 12:12 h light–dark cycle (lights on at 0630 h) and all experimental manipulations were performed during the light cycle (between 0900–1600 h). Mice were housed in same-sex groups of 2-5 mice per cage with standard laboratory chow and water available *ad libitum*. Male mice heterozygous for the *Hnrnph1* deletion were generated on a C57BL/6J background and were crossed at each generation to female wild-type C57BL/6J mice that were freshly ordered from The Jackson Laboratory (Bar Harbor, ME USA) to prevent genetic drift of the isogenic C57BL/6J background. Offspring included approximately 50% WT mice and 50% H1^+/-^ mice.

### Genotyping

Genotyping was conducted as previously described (Yazdani et al. 2015). We designed forward (GTTTTCTCAGACGCGTTCCT) and reverse (ACTGACAACTCCCGCCTCA) primers upstream and downstream of the TALENs binding domain (used to generate the deletion) within exon 4 of *Hnrnph1*. DreamTaq Green PCR Mastermix (ThermoScientific) was used in PCR amplification a 204 bp region containing the deletion site. PCR products were incubated overnight at 60 °C with either BstNI restriction enzyme (New England Biolabs) or a control buffer solution without enzyme. Gel electrophoresis (2% agarose gel with 0.5 μg/mL ethidium bromide) was performed on enzyme-treated and control samples for band visualization under UV light. The 204 bp PCR amplicon contained 2 BstNI restriction sites that flanked the cut sites of the TALENs *Fok*I nuclease. Mice carrying a heterozygous *Hnrnph1* deletion displayed 2 bands on the gel, while WT mice showed only a single band.

### Surgery and Histology

Age-matched female and male H1^+/-^ and WT mice were 50-70 days old at the time of surgery. Mice were anesthetized with 1-4% isoflurane and underwent stereotaxic surgery to implant a bipolar stimulating electrode (Plastics1, Roanoke, VA; MS308) that was aimed at the medial forebrain bundle (**MFB**) at the level of the lateral hypothalamus (−1.7, 1.0, −5.0 mm; relative to Bregma) (Paxinos and Franklin 2004). The analgesic meloxicam (5.0 mg/kg, s.c.) was administered on the day of surgery and on the three days following the surgery. The mice recovered for one week prior to beginning ICSS training. Following testing, mice were anesthetized with isoflurane and were transcardially perfused with 0.9% saline followed by 4.0% paraformaldehyde in 0.1 M PBS to fix brain tissue. Brains were later sectioned with a cryostat (40 µm), stained with 1% cresyl violet (Nissl stain), and imaged under a light microscope (4x) to confirm accurate electrode placement in the MFB. Placements were assessed blinded to the behavioral data, genotype, or sex. All mice that completed the behavioral testing were included in the analysis.

### ICSS Training

#### FR1 training

Mice were trained on a fixed-ratio 1 (**FR1**) schedule to respond for stimulation via a wheel manipulandum (ENV-113AM; Med Associates, St. Albans VT USA) within an operant chamber (15.24 x 13.34 x 12.7 cm; ENV-307A-CT; Med Associates) as previously described (Fish et al. 2012, Carlezon and Chartoff 2007). Operant chambers were placed inside closed sound-attenuating cubicles (ENV-022V; Med Associates). For every ¼ turn of the response wheel, a 500 ms train of square-wave, cathodal current (100-ms pulses) was delivered at a constant frequency of 142 Hz through a stimulator (PHM-150B/2; Med Associates). Each stimulation was followed by a 500 ms time-out period during which responses were counted but not reinforced by stimulation. The current intensity was adjusted for each subject to the lowest value that produced > 500 responses during a 45-min period across at least 3 consecutive training sessions (−50 to −140 uA). The minimum effective current for each subject was held constant throughout the rest of the study. All behavioral procedures were performed using Med-PC V software (Med Associates).

#### Rate-frequency training

This procedure has been described previously (Carlezon and Chartoff, 2007). Mice were introduced to stimulation delivered at their respective minimum effective currents over a series of 15 descending frequencies (log_0.05_ steps, 142-28 Hz). Each frequency trial consisted of 5 s of non-contingent priming stimulation, a 50-s response period during which time, responses were rewarded with 500 ms stimulation, and a 5-s time-out period. Each response-contingent stimulation was followed by a 500 ms timeout. This sequence was repeated for all 15 frequencies (1 pass = 15 min), and the procedure was repeated two more times in the same session (three total passes = 45 min). The number of responses (including those emitted during timeout periods) and stimulations delivered were recorded for each trial. Training was repeated until mice consistently responded at maximum rates during the highest frequency trials and did not respond during the lowest frequency trials. Subjects were required to demonstrate less than 15% between-day variability in mean response threshold for at least 3 consecutive training days prior to testing with MA.

### ICSS Testing

Mice were weighed and assessed for baseline, drug-naïve ICSS responding via a rate-frequency procedure that was identical to what was used during training (see above). Upon completion of the 45-min baseline procedure, mice were removed from the operant chambers, injected with MA (0.5, 1.0, 2.0, 4.0 mg/kg; i.p.), and placed individually into new, clean cages with fresh bedding. Ten min post-MA injection, mice were placed back into the operant chambers and underwent the same protocol used during baseline assessment (3 passes of 15 min, 45 min total). Testing was performed every other day and all mice received escalating doses of MA using a within-subjects design. On testing days, videos of both baseline and post-MA sessions were recorded for later behavioral tracking and analysis using ANY-maze V software (Stoelting Co., Wood Dale, IL).

### ANALYSIS

#### Minimum effective current

Minimum effective currents were analyzed using 2-way analysis of variance (**ANOVA**) test with Genotype and Sex as between-subject factors.

#### M50

M50 values reflect the frequency at which a mouse responds at half of its maximum response rate for a given pass; this measure is analogous to an effective concentration (**EC**) 50 value of a pharmacological dose-response curve. Just as a decrease in EC50 value indicates increased drug potency, a decrease in M50 value indicates *heightened sensitivity* to the reward-eliciting properties of MFB stimulation. Detailed descriptions of this approach are published (Miliaressis et al. 1986; Carlezon and Chartoff 2007). M50 values for each pass were determined using a custom-built analysis program and were averaged across all three passes in a baseline or MA session.

We analyzed both raw M50 data and M50 data normalized to same-day baseline values. For the raw data analysis, the mean baseline M50 score across all baseline sessions was used (prior to the 0.5, 1.0, 2.0, and 4.0 mg/kg doses) and compared to post-MA M50 values following each dose. This analytical approach allowed us to detect shifts in M50 values relative to baseline, indicating MA-induced changes in reward sensitivity. As M50 values are frequently presented as normalized values (% baseline M50 = post-drug M50 ÷ baseline M50 x 100), we additionally analyzed post-MA values that were normalized to same-day baseline M50 values. This method adjusts for between-day baseline variability and permits comparison of normalized effects of MA at each dose. Raw M50 values were analyzed via three-way repeated measures (**RM**-) ANOVAs with Genotype and Sex as between-subjects factors and MA Dose (baseline, 0.5, 1.0, 2.0, 4.0 mg/kg) as the repeated measure. Similarly, percent baseline M50 values were analyzed via three-way repeated measures (**RM**-) ANOVAs with Genotype and Sex as between-subjects factors and MA Dose (0.5, 1.0, 2.0, 4.0 mg/kg) as the repeated measure. We additionally performed RM-ANOVAs on baseline M50 values collected prior to each MA dose to ensure there were no significant shifts across testing days, including Genotype and Sex as between-subjects factors.

Significant interactions were further pursued by deconstructing along the appropriate factor (Genotype or Sex) and running two-way RM-ANOVAs with either Genotype or Sex as the between-subjects factor and MA Dose as the repeated measure. Post-hoc differences were determined via two-tailed Student’s t-tests with significance set to α=0.05 and Bonferroni correction used to adjust for multiple comparisons. All within-subjects pairwise comparisons were made to baseline values (for the raw data analysis) or 0.5 mg/kg percent baseline values (for the normalized analysis).

#### Maximum response rates

In addition to measuring changes in sensitivity to MFB stimulation (via M50 values), we also assessed changes in operant response rates. Relative to baseline rates, differences in maximum response rates and total responses can reflect the effect of MA on both reward and operant performance. Average maximum response rates during any frequency trial and total responses were determined for each baseline and MA session. Like the M50 data, we analyzed both raw and normalized data, as well as baseline values across each testing day. Similarly, the baseline value that was compared to raw post-MA value was determined by averaging across all baseline sessions on that particular day (prior to every MA dose). Percent baseline values reflect changes from same-day baseline maximum response rates or total responses. RM-ANOVAs and post-hoc analyses of maximum response rate and total response data were run in the same manner as the M50 analysis described above.

#### Concomitant behavior

Behavioral data were collected using custom tracking protocols in ANY-maze V (Stoelting Inc. Wood Dale, IL, USA). The average distance to the response wheel was determined by setting a point of interest at the central-most position of the response wheel using ANY-maze. Both raw and normalized data were analyzed. The average baseline distance was determined across all baseline sessions and used in the raw post-MA for each dose. Percent baseline values were calculated using same-day baseline values. RM-ANOVAs and post-hoc analyses of wheel distance data were run in the same manner as the M50 and operant responding data described above.

## RESULTS

### Electrode placement

We analyzed data from a total of 34 mice. Histological analysis confirmed accurate electrode placement in the MFB or in a proximity to these fibers sufficient to produce activation (**Fig. 1a-b**) in all mice that completed MA testing (n=34; 14 H1 WT (5 females (**F**); 9 males (**M**)) and 20 H1^+/-^ (10 F, 10 M)). Average electrode placements were −1.89 ± 0.04 (mean ± SEM) AP, 1.00 ± 0.03 ML, and −5.05 ± 0.03 DV (mm relative to Bregma). A total of 6 out of 40 surgerized mice were omitted from all analyses because they either failed to meet training criteria or because the electrode detached prior to completing the study.

**Fig 1.**
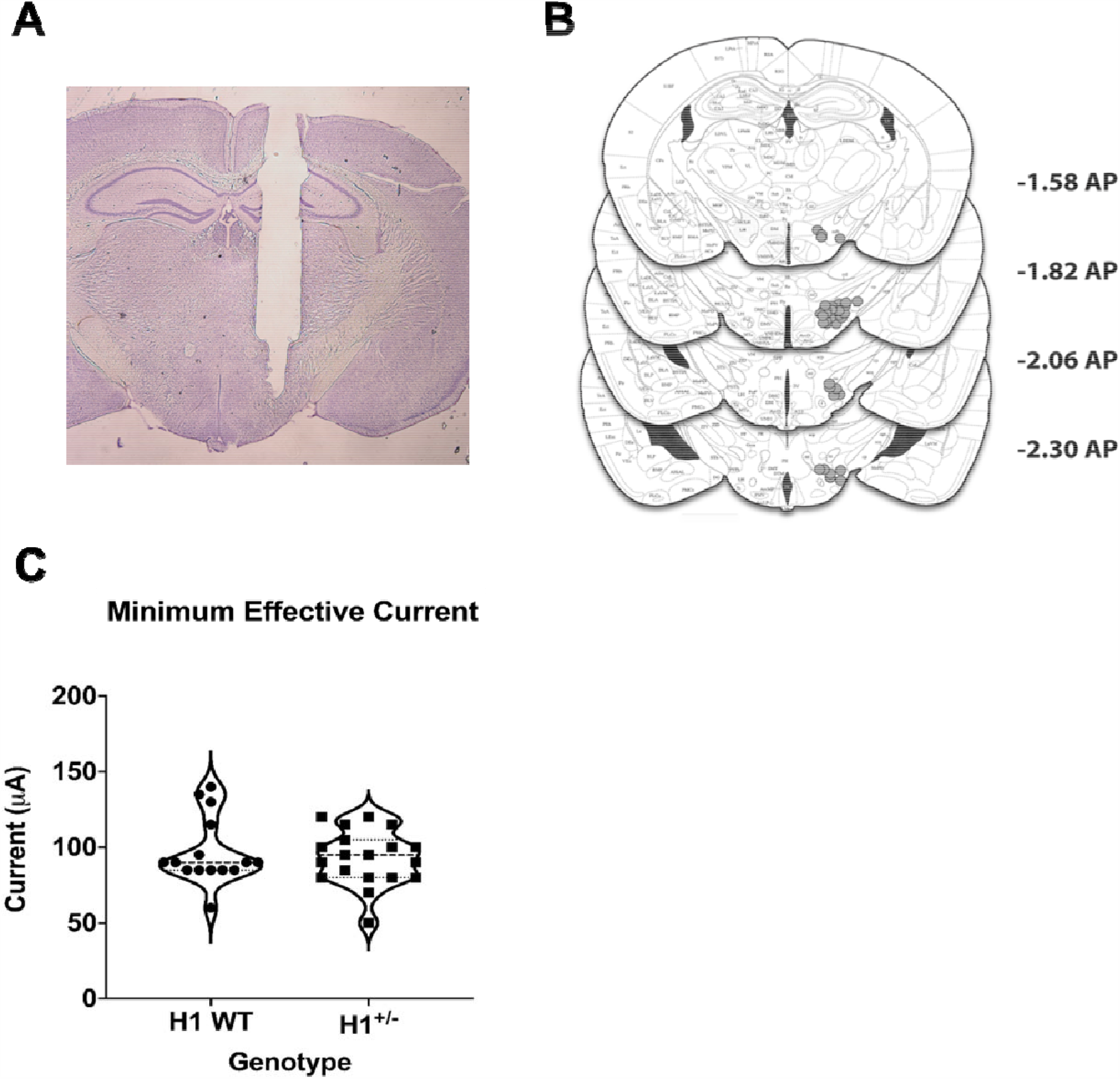
Electrode placements in the medial forebrain bundle (MFB) and minimum effective currents (MECs) in H1 WT (n=14) and H1^+/-^ (n=20) mice. **a**: Representative photomicrograph of an electrode tract and terminus in the MFB. **b**: Schematic of all electrode placements with anterior/posterior (AP) coordinates of each coronal plane listed on the right (relative to Bregma; Paxinos and Franklin 2004). **c**: Violin plot of minimum effective current distribution. A 2-way ANOVA revealed no significant effect of Genotype or Sex on MECs and no interaction (all p’s>0.28)

### Minimum Effective Current (MEC)

The average MEC was −95.0 ± −3.4 (SEM) µA. There was no significant effect of H1 Genotype or Sex and no interaction (**Fig. 1c**).

### Operant Responding

No significant dose-dependent effect of MA was observed on the total number of unnormalized, raw responses (**Fig. 2a**). However, when adjusting for day-to-day baseline response values, there was a significant effect of MA Dose on the normalized total number of responses (**Fig. 2b**), with a dose-dependent increase that was significant following the 2.0 mg/kg and 4.0 mg/kg MA doses relative to the 0.5 mg/kg MA dose.

**Fig 2.**
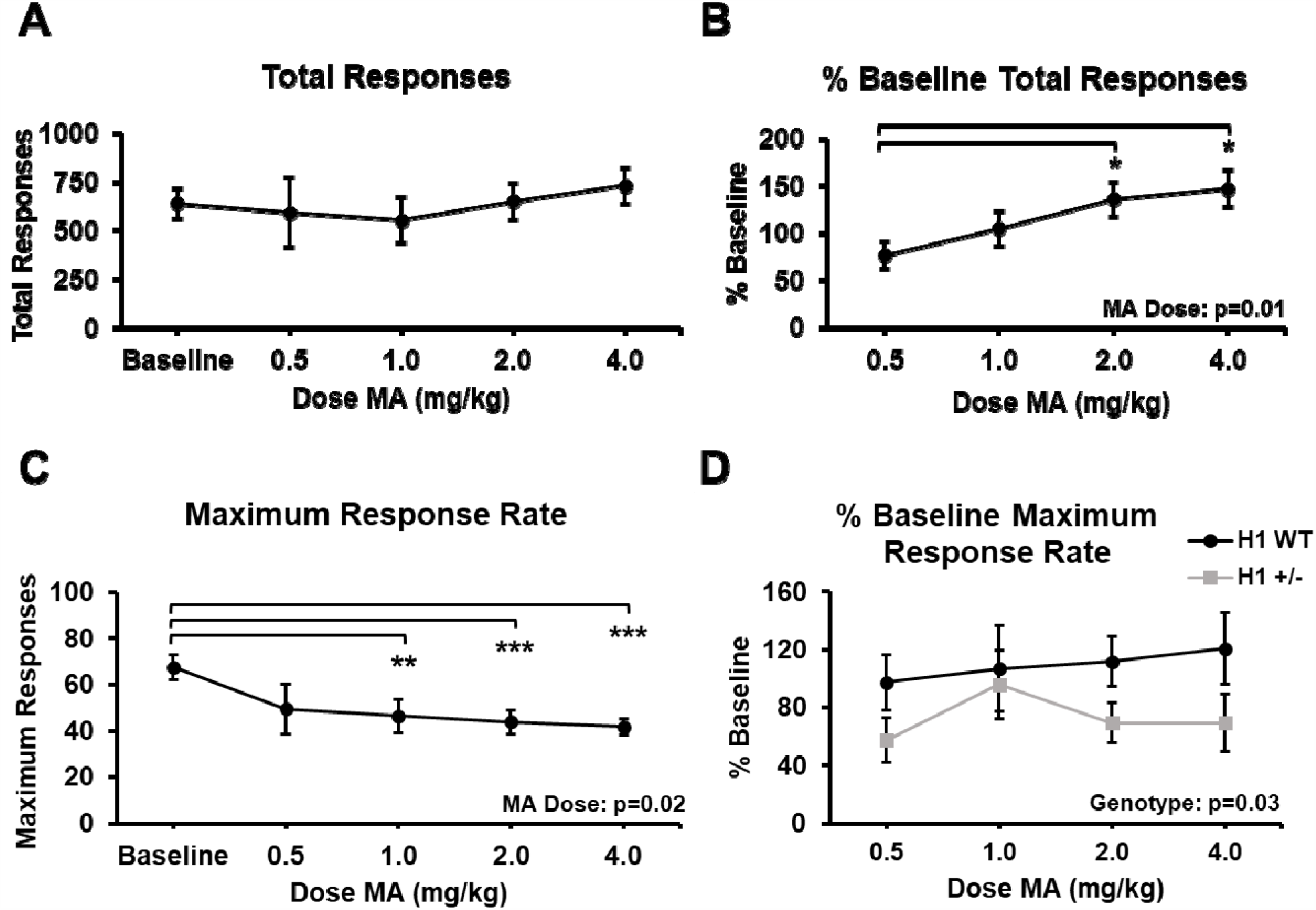
MA treatment increases normalized total responses and decreases raw maximum response rates. **a:** Raw total ICSS responses at baseline and following increasing MA doses (n=34; 15 F,19 M). There was no effect of MA Dose or Genotype (both p>0.66). **b:** Total responses normalized to same-day baseline values. There was a main effect of MA Dose (F_3,90_=3.83, p=0.01). Normalized total responses were significantly higher relative to 0.5 mg/kg values following both 2.0 and 4.0 mg/kg MA (Student’s paired t-tests; both Bonferroni adj. p<0.05). **c:** Raw maximum response rates at baseline and following increasing MA doses. There was a main effect of MA Dose (F_4,120_=2.79, p=0.02) that was explained by a significant reduction in response rate compared to baseline following 1.0 (p<0.01), 2.0 (p<0.001), and 4.0 mg/kg (p<0.001) MA. **d:** Maximum response rates normalized to same-day baseline values following increasing MA doses [n=20 H1^+/-^ (10 F, 10 M) and n=14 H1 WT (5 F, 9 M)]. There was a main effect of H1 Genotype (F_1,30_=5.01, p=0.03) interactions, indicating an overall decrease in the rate, irrespective of MA dose. Significant main but no effects indicated within the figure panels with their respective p values. Pairwise comparisons were performed using paired Student’s t-tests with Bonferroni correction against raw baseline or 0.5 mg/kg % baseline Significance markers: *=p<0.05, **=p<0.01, ***=p<0.001 values.

For raw maximum response rates, we observed significant decreases at 1.0, 2.0, and 4.0 mg/kg MA when compared to the average baseline rates (**Fig. 2c**). For normalized maximum response rates, there was a significant effect of H1 Genotype was driven by normalized rates in H1 WT mice that were an average of 36.20% higher than those of H1^+/-^ mice (Confidence Interval (**CI**) for Mean Difference (H1 WT – H1^+/-^): [3.15, 69.24]; p=0.03; **Fig. 2d**). Maximum response rates and total responses recorded during baseline sessions were not significantly different across test days and were not significantly affected by Genotype or Sex (all p’s>0.08; **Suppl. Fig. 1b-c)**.

### M50

For raw M50 values, there was a main effect of MA Dose, with significant decreases observed following 2.0 mg/kg and 4.0 mg/kg (**Fig. 3a**), indicating an overall increase in sensitivity to brain stimulation reward relative to baseline conditions (Miliaressis et al. 1986; Carlezon and Chartoff 2007). For normalized M50 values, there was a significant Genotype x Sex interaction (**Fig. 3b**) that was partially driven by values that were, on average, 15.38% lower in H1^+/-^ compared to WT females (95% CI for Mean Difference (H1 WT F – H1^+/-^ F): [2.67, 28.09]; p=0.02), suggesting an overall greater reward sensitivity in H1^+/-^ females versus WT females in response to MA, irrespective of MA Dose. In addition, we found a significant decrease in M50 values in females of both genotypes from 0.5 mg/kg MA to 2.0 mg/kg MA (**Fig. 3c**), suggesting enhanced reward sensitivity induced by the higher MA dose in females. For males, there was no Dose or Genotype effect or interaction in normalized M50 values (**Fig. 3d**). We also note that there were no effects of Genotype, Sex, or Dose on baseline M50 values collected prior to each MA session (**Suppl. Fig. 1a**).

**Fig 3.**
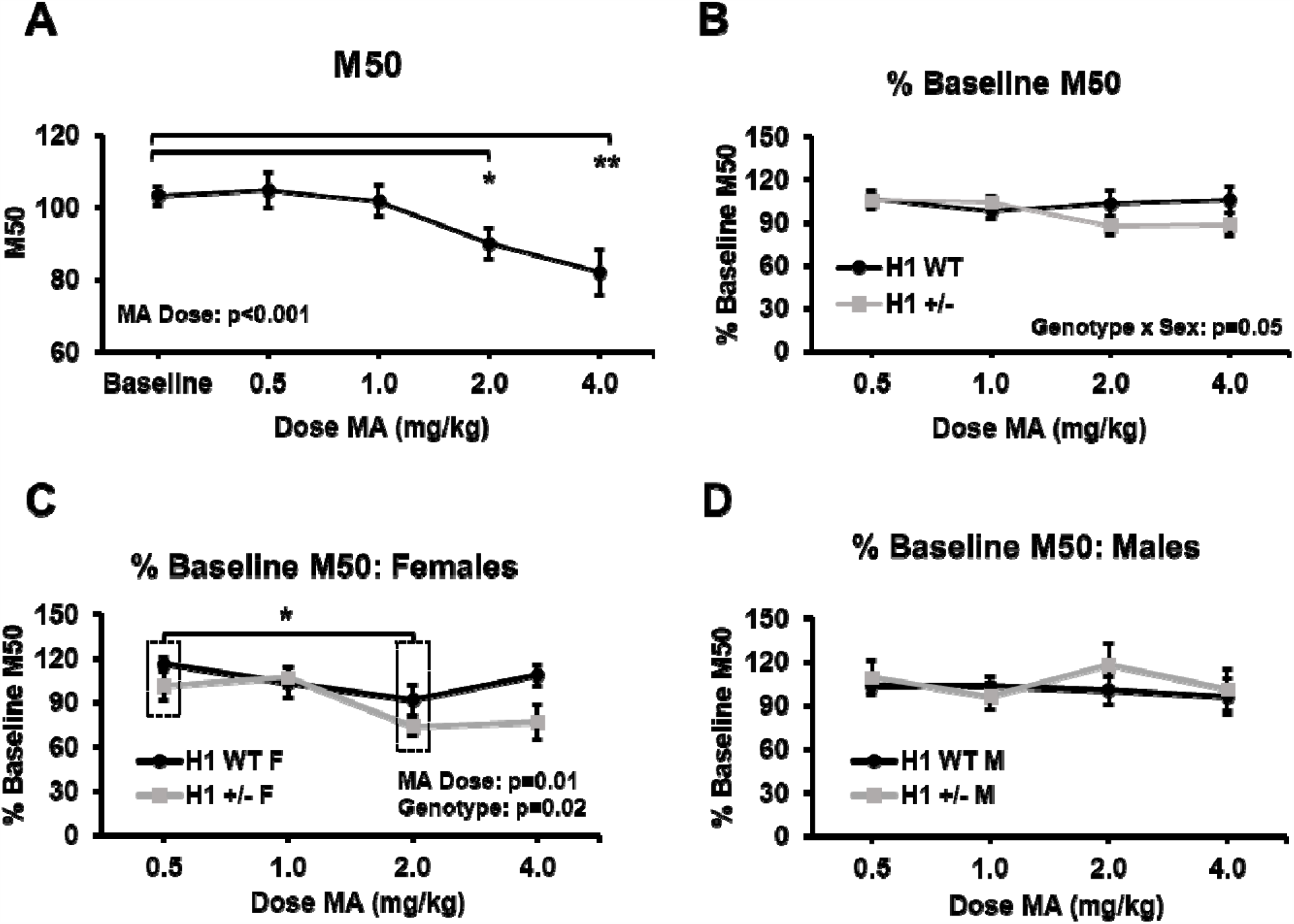
Raw M50 values are significantly reduced following increasing MA doses and normalized M50 values vary in a genotype-, dose-, and sex-dependent manner. **a:** Raw M50 values at baseline and following increases doses of MA (n=34). There was a significant main effect of MA Dose (F_4,120_=6.82, p<0.001) and M50 values were significantly lower than baseline values following both 2.0 (p<0.05) and 4.0 (p<0.01) mg/kg MA. **b:** M50 values normalized to baseline values following each dose MA [n=20 H1^+/-^ (10 F, 10 M) and n= 14 H1 WT (5 F, 9 M)]. There was a near-significant Genotype x Sex interaction (F_1,30_=4.12, p=0.05). These data are shown for the individual sexes in panels **c** (females) and **d** (males). **c**: In females (n=5 H1 WT, 10 H1^+/-^), there were significant main effects of MA Dose (F_3,42_=3.50, p=0.01) and H1 Genotype (F_1,14_=6.73, p=0.02). Across all doses, H1 WT females had normalized M50 values that were 15.38% higher than H1 WT females (95% Confidence Interval for Mean Difference (H1 WT–H1^+/-^): [2.67, 28.09]; p=0.02). Percent baseline values were significantly lower following the 2.0 mg/kg dose relative to the 0.5 mg/kg dose for females of both genotypes (p=0.01; Genotype-independent effect indicated by dashed boxes in panel **c**). **d**: In males n=9 H1 WT,10 H1^+/-^), there were no significant main effects or interactions on normalized M50 values (all p>0.52). All data are presented as the mean ± S.E.M. P-values for significant main effects and interactions are indicated within the respective figure panels. Pairwise comparisons were performed using paired Student’s t-tests with Bonferroni corrections for raw baseline or 0.5 mg/kg % baseline values. Significance markers: *=p<0.05, **=p<0.01, ***=p<0.001

### Concomitant Behavior: Distance to the Response Wheel

We collected data from a total of 27 mice (11 H1 WT (5 F, 6 M) and 16 H1^+/-^ (7 F, 9 M)). Seven mice were excluded from this analysis due to errors with the recording equipment or the inability of the tracking software to track the mice. We found that mice were generally located closer to the wheel following increasing doses of MA, and this effect was significant at the 4.0 mg/kg MA dose relative to baseline distance (**Fig. 4a**). There was also a significant interaction between Genotype and Sex. When breaking down the data by Sex, there was a main effect of MA Dose in both sexes (both p’s<0.005), but no main effect or interaction with Genotype (all p>0.12). Comparatively, breaking down the data by Genotype identified a significant effect of Dose in H1^+/-^ mice, with both sexes being located closer to the response wheel following 4.0 mg/kg MA than they were at baseline. There was also a main effect of Sex that was specific to H1^+/-^ mice, as H1^+/-^ males tended to be closer to the response wheel than H1^+/-^ females regardless of MA dose. When considering distance relative to baseline values, mice were located significantly closer to the response wheel following 2.0 and 4.0 mg/kg of MA than they were following 0.5 mg/kg MA (**Fig. 4b**), suggesting dose-dependence for this phenotype. Interestingly, we detected a significant Sex x Genotype interaction for wheel distance during baseline sessions (p=0.008; **Suppl. Fig. 1d**) that was ultimately explained by female H1^+/-^ mice being located closer to the wheel than female H1 WT mice when averaged across all baseline sessions (p=0.03; **Suppl. Fig. 1e**). There were no significant effects in the males (**Suppl. Fig. 1f**).

**Fig 4.**
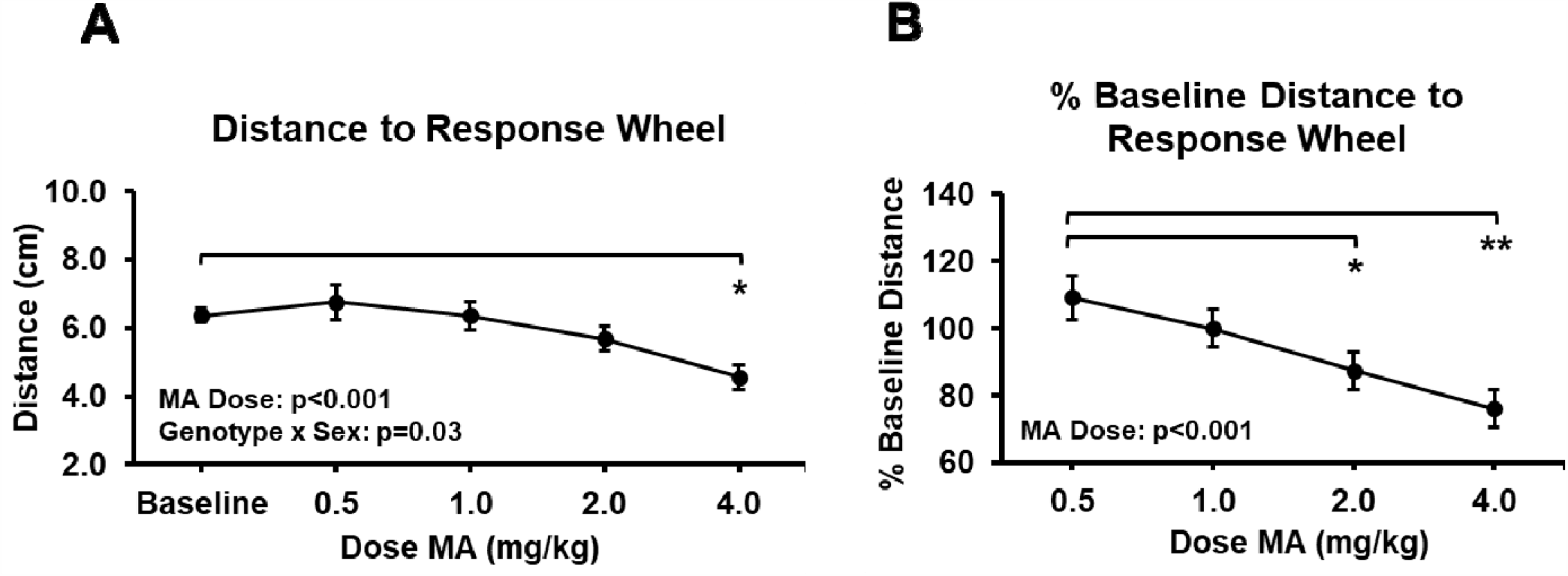
MA significantly decreases average distance to the ICSS response wheel. **a:** Raw data reflect the average distance to the response wheel at baseline and for each dose MA (n=27). There was a significant main effect of MA Dose (F_4,92_=8.32, p<0.001) as well as a significant Genotype x Sex interaction (F_1,23_=5.04, p=0.03; n=16 H1^+/-^ (7 F, 9 M) and n=11 H1 WT (5 F, 6 M)). Pairwise comparisons to baseline distance revealed mice were located significantly closer to the response wheel following 4.0 mg/kg MA (p<0.05). **b:** Distance to the response wheel is normalized to same-day baseline values for each MA dose. There was a significant main effect of MA Dose on normalized distance (F_3,69_=6.71, p<0.001), with significantly lower values following 2.0 and 4.0 mg/kg MA relative to 0.5 mg/kg values (2.0 mg/kg vs. 0.5 mg/kg: p<0.05; 4.0 mg/kg vs. 0.5 mg/kg: p<0.01). All data are presented as the mean ± S.E.M. Significant main effects and interactions indicated within Fig. panels with their respective p values. Pairwise comparisons were performed using paired Student’s t-tests with Bonferroni correction against raw baseline or 0.5 mg/kg % baseline values. Significance markers: *=p<0.05, **=p<0.01

## DISCUSSION

Previous work has consistently shown that monoamine releasers such as MA and amphetamine facilitate ICSS reward as evidenced by elevated response rates and decreased reward thresholds relative to baseline or vehicle conditions (Bauer et al. 2013; Esposito et al. 1980; Negus and Miller 2014; Robinson et al. 2012). We demonstrate that MA facilitates ICSS by dose-dependently increasing total responses relative to baseline conditions (**Fig. 2b**) and dose-dependently reducing raw M50 values for reward sensitivity (**Fig. 3a**). We also identified concomitant behaviors showing a dose-dependent increase in proximity of the mice to the response wheel following increasing doses of MA (**Fig. 4a-b**). Together, these overall effects of MA validate our ICSS procedure in examining brain stimulation reward and MA facilitation.

Based on our prior observation of reduced MA-induced locomotor activity, dopamine release, reward, and reinforcement in H1^+/-^ mice, we hypothesized that H1^+/-^ mice would show changes in ICSS and MA facilitation of brain stimulation reward. No differences were observed in MEC (**Fig. 1**) or drug-naïve M50 values (**Suppl. Fig. 1a**), indicating similar sensitivity to brain stimulation reward in H1^+/-^ mice prior to MA exposure. This negative finding aligns with our previous observations in drug-naïve H1^+/-^ mice, as microdialysis studies revealed no effect of H1^+/-^ on total tissue levels or extracellular levels of dopamine in the nucleus accumbens at baseline (Ruan et al. 2020b).

In response to MA, H1^+/-^ mice showed lower normalized maximum response rates, irrespective of MA Dose (**Fig. 2d**), providing one piece of evidence to support our hypothesis of blunted MA facilitation of brain stimulation reward in H1^+/-^ mice. We also note that these lower maximum response rates are unlikely to be explained by motor-related performance deficits since there was no effect of H1^+/-^ on total responses following MA administration (**Fig. 2a-b**).

H1^+/-^ females, on average, showed overall lower normalized M50 values than H1 WT females across MA doses as indicated by a significant Genotype effect (**Fig. 3c**), providing evidence for increased sensitivity to MA-induced reward facilitation. This increased sensitivity was unexpected, given that H1^+/-^ mice (irrespective of sex) showed blunted MA-induced DA extracellular levels in the NAc compared to wild-type (**WT**) littermates following intraperitoneal (i.p.) injections of MA (0.5 and 2.0 mg/kg) (Ruan et al. 2020b). Thus, the combination of MA stimulation and ICSS could induce a distinct profile of neurochemical release and/or neuronal activation within the mesocorticolimbic circuitry in H1^+/-^ females compared to H1 WT females.

It is not clear what mechanisms underlie the sex effects observed in this study (**Fig. 3b-d**) but could potentially be attributable to different pharmacokinetic and/or metabolic profiles between the sexes (Lominac et al. 2014) or to differences in pharmacodynamics of MA-induced regulation of DAT or VMAT that can affect the time course of changes in DA neurotransmission (Dluzen et al. 2008; Dluzen & McDermott 2008; Milesi-Hallé et al. 2005). More broadly, the effects of H1^+/-^ on post-MA ICSS responding could be mediated by differences in the pharmacodynamics of DA release and signaling in the NAc (Otani et al. 2008; P Chan et al. 1994; Albers and Sonsalla 1995; Bousman et al. 2009b). Furthermore, we previously found no genotypic difference in brain concentration of MA or its metabolite at 30 min post-MA (Ruan et al. 2020b). Nevertheless, a full time-course of brain and plasma concentrations at different doses would more fully address the question of potential sex-dependent pharmacokinetic differences and MA bioavailability in H1^+/-^ mice.

A novel component of this study comprised the identification of a concomitant behavior during ICSS testing. We observed a MA dose-dependent decrease in both normalized and raw average distance to the operant response wheel (**Fig. 4a-b**), which likely represents both approach and consummatory behaviors since an increase in ICSS responding necessitates an increase in time spent near the response wheel. Reinforcement-induced place preference for the microenvironment associated with MA-modulated reward potentiation could also increase proximity for the ICSS microenvironment.

Despite the significant decrease in MA-induced locomotor activity that we have repeatedly observed following 2.0 mg/kg (i.p.) in H1^+/-^ mice (Yazdani et al. 2015; Ruan et al. 2020b), we did not detect any genotypic difference in MA-induced locomotor activity during ICSS testing sessions. This discrepancy is likely explained by the much smaller size of the operant ICSS chambers compared to the open field and by the restricted movement associated with being attached to the stimulation apparatus. In support, the reduction in MA-induced locomotor activity in H1^+/-^ mice was much more robust in an undivided open field arena (Yazdani et al. 2015) compared to the enclosed MA side of the CPP chamber (Ruan et al. 2020b).

Our sample size was based on a prior power analysis of previous sample size estimate for detecting genotypic effects on MA-induced locomotor activity (Cohen’s d = 0.9; required n=16 to achieve 80% power; p<0.05) (Ruan et al., 2020b). Another limitation is that we administered increasing doses of MA to all mice, preventing our ability to dissociate the effect of prior drug exposure with the effect of the current MA dose. Future studies that employ larger sample sizes based on the effect sizes of the present study and that employ a between-subjects design for each dose will be necessary to ensure the reliability and generalizability of these findings.

In conclusion, irrespective of genotype, we demonstrate dose-dependent MA-induced facilitation of operant brain stimulation reward and dose-dependent induction of approach behaviors toward the microenvironment associated with this reward. We also present evidence for MA-induced modulation of brain stimulation reward in H1^+/-^ mice in the absence of any Genotype effect on baseline ICSS responding, including no effect on drug-naïve minimally effective currents or M50 values. The data suggest H1 mutation induces selective effects on MA-induced facilitation of brain stimulation reward and no detectable effects on basal innate reward processing following MFB stimulation, which aligns with our previous findings indicating selective effects on MA-induced, but not basal, neurobehavioral phenotypes. We identified sex- and MA dose-dependent differences in modulation of brain stimulation reward in H1^+/-^ mice that could behaviorally reflect an interaction of ICSS and altered dynamics and magnitude of MA-induced extracellular dopamine levels (Ruan et al. 2020b). Ongoing studies are aimed at deciphering the molecular mechanisms by which H1^+/-^ robustly decreases MA-induced dopamine release and behavior in H1^+/-^ mice.

## ACKNOWLEDGEMENTS AND DISCLOSURES

This work was funded by R01DA039168 (C.D.B.), U01DA050243 (C.D.B.), NIGMS T32 Biomolecular Pharmacology Training Grant GM008541 (K.N.B.), and Boston University’s Transformative Training Program in Addiction Science (TTPAS Burroughs Wellcome Fund: #1011479; K.N.B.). We thank Julia L. Scotellaro for assistance with genotyping, histology, and behavioral training. Dr. Carlezon discloses that he has served as a paid consultant for Psy Therapeutics within the past 2 years.

## SUPP. FIGURE 1

**Suppl. Fig. 1.**
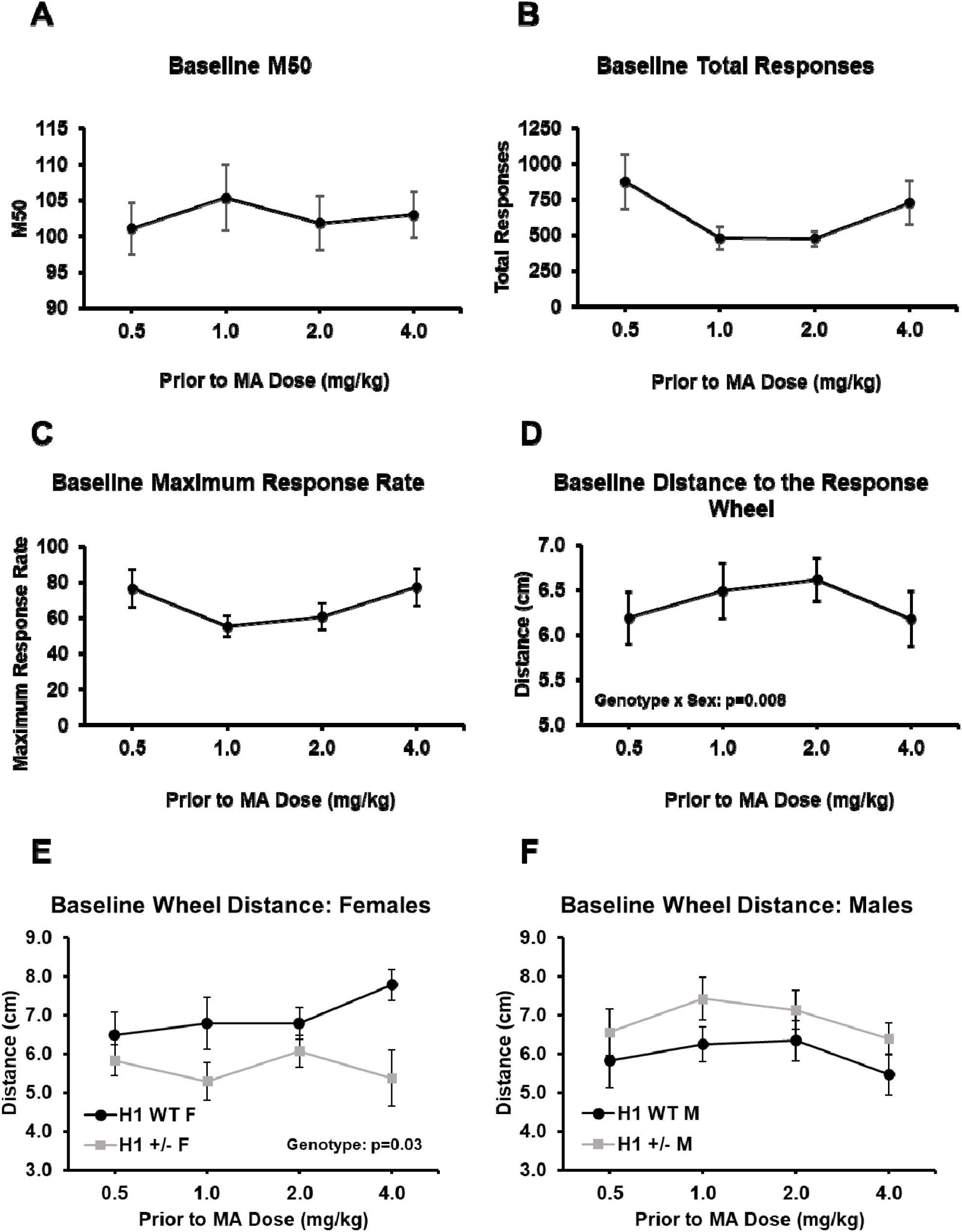
H1^+/-^ does not affect baseline M50 values, total responses, or maximum response rates but decreases distance to response wheel in females. Data collected from baseline sessions immediately preceding the corresponding MA dose including M50 values (**a**), total responses (**b**), maximum response rates (**c**), and distance to the response wheel (**d-f**). **a:** There were no significant effects or interactions of Genotype, Sex, or Dose on M50 values at baseline (n=34; all p>0.19). Similarly, there were no significant effects or interactions on total responses (**b**; all p>0.09) or maximum responses (**c**; all p>0.08). **d**: Distance to the response wheel across baseline sessions revealed a significant Genotype x Sex interaction (n=16 H1^+/-^ (7 F, 9 M) and n=11 H1 WT (5 F, 6 M); F_1,23_=8.54, p=0.008). Breaking down the data by Sex revealed a significant effect of Genotype in the females (**e**; F_1,10_=6.11, p=0.03) and no significant main effects or interactions in the males (**f**; all p>0.11). Across all doses, H1^+/-^ females were located 1.32 cm closer to the response wheel than H1 WT females during baseline sessions (p=0.03).

