## Supplemental Figure 1 for "Intracranial self-stimulation and concomitant behaviors following systemic methamphetamine administration in *Hnrnph1* mutant mice"

**SUPP. FIGURE 1**

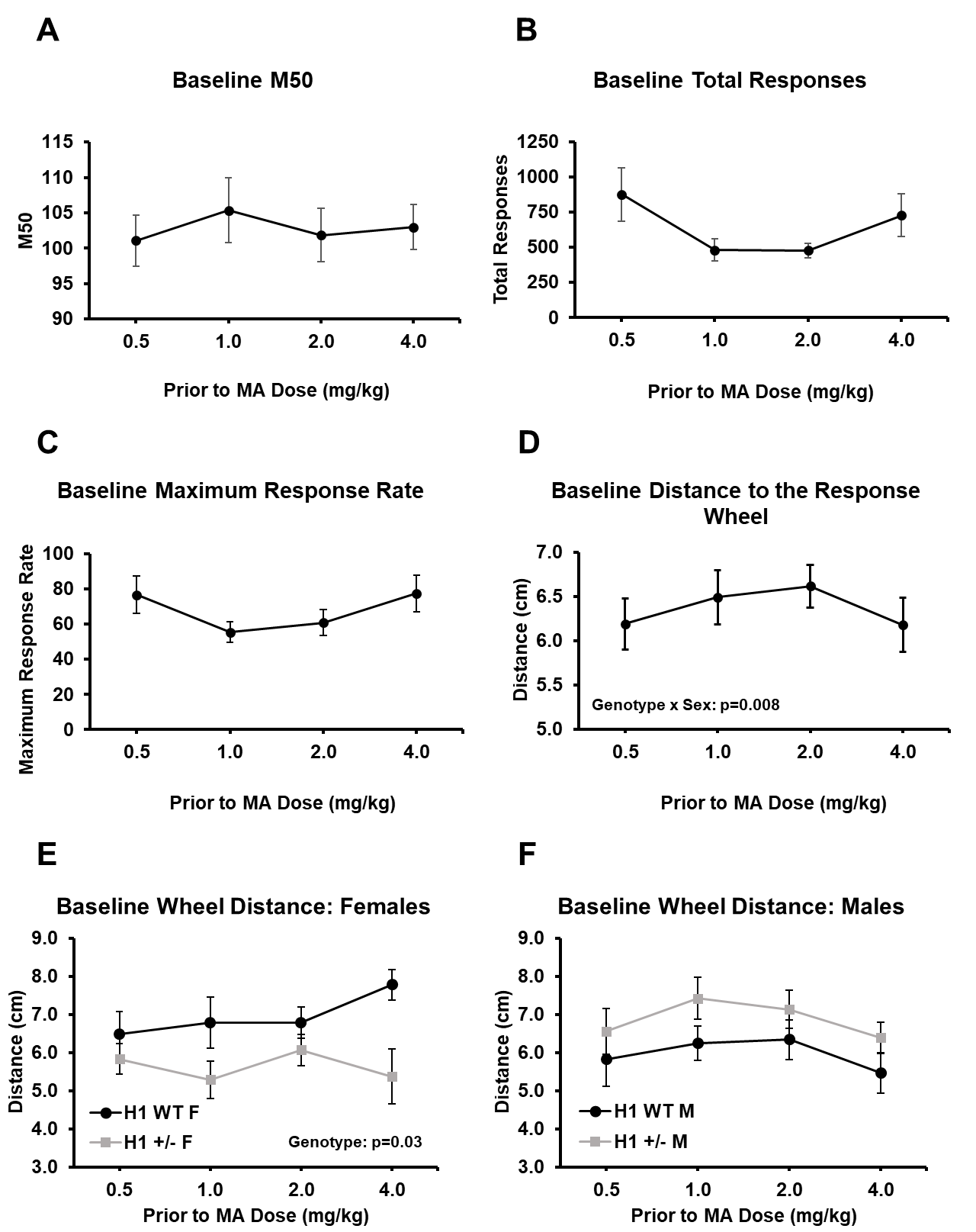

**Suppl. Fig. 1. H1^+/-^ does not affect baseline M50 values, total responses, or maximum response rates but decreases distance to response wheel in females.** Data collected from baseline sessions immediately preceding the corresponding MA dose including M50 values (**a**), total responses (**b**), maximum response rates (**c**), and distance to the response wheel (**d-f**). **a:** There were no significant effects or interactions of Genotype, Sex, or Dose on M50 values at baseline (n=34; all p>0.19). Similarly, there were no significant effects or interactions on total responses (**b**; all p>0.09) or maximum responses (**c**; all p>0.08). **d**: Distance to the response wheel across baseline sessions revealed a significant Genotype x Sex interaction (n=16 H1^+/-^ (7 F, 9 M) and n=11 H1 WT (5 F, 6 M); F_1,23_=8.54, p=0.008). Breaking down the data by Sex revealed a significant effect of Genotype in the females (**e**; F_1,10_=6.11, p=0.03) and no significant main effects or interactions in the males (**f**; all p>0.11). Across all doses, H1^+/-^ females were located 1.32 cm closer to the response wheel than H1 WT females during baseline sessions (p=0.03).
